# Genetic variation for tolerance to the downy mildew pathogen *Peronospora variabilis* in genetic resources of quinoa (*Chenopodium quinoa*)

**DOI:** 10.1101/2020.08.19.257535

**Authors:** Carla Colque-Little, Miguel Correa Abondano, Ole Søgard Lund, Daniel Buchvaldt Amby, Hans-Peter Piepho, Christian Andreasen, Sandra Schmöckel, Karl Schmid

**Affiliations:** Institute of Plant Breeding, Department of Plant and Environmental Sciences, University of Copenhagen, Denmark; Institute of Plant Breeding, Seed Science and Population Genetics, University of Hohenheim, Stuttgart, Germany; Institute of Crop Science, University of Hohenheim, Stuttgart, Germany

**Keywords:** *Chenopodium quinoa*, *Chenopodium album*, *Peronospora variabilis*, downy mildew, phenotyping, linear mixed models, quantitative resistance, plant genetic resources

## Abstract

**Background:** Quinoa (*Chenopodium quinoa* Willd.) is an ancient grain crop that is tolerant to abiotic stress and has favorable nutritional properties. Downy mildew is the main disease of quinoa and is caused by infections of the biotrophic oomycete *Peronospora variabilis* Gaüm. Since the disease causes major yield losses, identifying sources of downy mildew tolerance in genetic resources and understanding its genetic basis are important goals in quinoa breeding.

**Results:** We infected 132 South American genotypes, three Danish cultivars and the weedy relative *C. album* with a single isolate of *P. variabilis* under greenhouse conditions and observed a large variation in disease traits like severity of infection, which ranged from 5% to 83%. Linear mixed models revealed a significant effect of genotypes on disease traits with high heritabilities (0.72 to 0.81). Factors like altitude at site of origin or seed saponin content did not correlate with mildew tolerance, but stomatal width was weakly correlated with severity of infection. Despite the strong genotypic effects on mildew tolerance, genome-wide association mapping with 88 genotypes failed to identify significant marker-trait associations indicating a polygenic architecture of mildew tolerance.

**Conclusions:** The strong genetic effects on mildew tolerance allow to identify genetic resources, which are valuable sources of resistance in future quinoa breeding.

## Background

Quinoa (*Chenopodium quinoa* Willd.) is a grain crop that was domesticated in South America and cultivated from Chile to Southern Colombia for thousands of years (Mujica and Jacobsen, 2007). After the arrival of the Spanish, it was replaced by European crops in many regions (Gómez and Aguilar, 2016). More recently, quinoa has experienced renewed interest as alternative grain crop worldwide and became an important export commodity for countries like Bolivia, where its exports in 2014 were approximately 40,000 tons (Gandarillas *et al*., 2015a). The interest in quinoa results from its nutritional properties and tolerance to abiotic stresses such as high salinity, drought, and frost (Gómez and Aguilar, 2016; Zurita *et al*., 2014). The increasing demand for quinoa and successful cultivation outside its native range led to multiple breeding programs aimed at improving yield, resistance and adaptation to novel cultivation regions or climate change (Bazile *et al*., 2015, 2016; Murphy *et al*., 2018). One important biotic factor that limits cultivation is susceptibility to plant diseases. Downy mildew, the most important disease of quinoa, is caused by the biotrophic oomycete *Peronospora variabilis* Gaüm, previously known as *Peronospora farinosa* f.sp. *chenopodii* (Choi *et al*., 2010). It causes severe yield losses of up to 30 *−* 50% in tolerant cultivars, and to an almost complete yield loss of 99% in susceptible cultivars under conditions of high humidity and absence of chemical control measures (Danielsen and Munk, 2004). The disease is widely spread over continents where quinoa is cultivated and may have been spread by seeds that were contaminated with the pathogen (Danielsen *et al*., 2002; Testen *et al*., 2012; Choi *et al*., 2014; Khalifa and Thabet, 2018). *P. variabilis* also infects the closely related and widespread weed *C. album* (Danielsen and Lübeck, 2010; Kara *et al*., 2020) which may act as a secondary host. Because the disease is seedborne, tolerance to this pathogen is a critical trait in the development of new quinoa varieties (Testen *et al*., 2014).

Currently, very little is known about the physiological mechanisms involved in the *P. variabilis* - quinoa interaction, or about the genetic basis of downy mildew tolerance and the role other phenotypic traits in disease susceptibility. Previous studies for quinoa tolerance using greenhouse experiments, seedlings, detached leaves and field scorings primarily focused on quantitative measures by scoring disease symptoms (Ochoa *et al*., 1999; Bonifacio, 2003; Danielsen and Munk, 2004; Khalifa and Thabet, 2018). Response to mildew infection utilizes visual scoring of disease severity, which is the proportion of leaf tissue with lesions caused by the pathogen (Danielsen and Munk, 2004). Another measure is the extent of sporulation by the pathogen. It is measured with a detached leaf assay and the identification of spore bodies on leaf surfaces (Benlhabib *et al*., 2016). Reliable and efficient scoring of tolerance to downy mildew is a key component in the development of improved quinoa varieties.

The objectives of the present study were to investigate the variation of quinoa genotypes from its native range in South America (Bolivia, Peru, Ecuador, and Chile) in their response to inoculation with the downy mildew pathogen. We investigated the robustness of phenotypic scoring under controlled conditions and characterized the relationship of the disease traits severity of infection, sporulation and incidence with other phenotypic traits. These traits included size and density of leaf stomata because *P. variabilis* enters leaf tissues through the stomata (Kitz, 2008; Choi *et al*., 2010), and seed saponin content because saponin extracts have antifungal properties (Jacobsen, 2003; Tenorio *et al*., 2010). We estimated genetic variance components and heritability of the response to *P. variabilis* infection and conducted a genome-wide association study (GWAS) with whole genome sequences a subset of accessions to identify genomic regions with putative tolerance genes.

## Results

### High variation in mildew tolerance

In total, 132 genotypes (5 controls, 21 cultivars and 106 accessions) were successfully grown, inoculated with mildew, phenotyped and scored in three independent greenhouse experiments. Severity of infection ranged from 5.0% (*Chenopodium album*) to 83.0% (Accession G9) with a mean of 46.2%, whereas sporulation ranged from 0.2% (Variety Puno) to 83.6% (Cultivar CV21) with a mean of 42.6%. Incidence of infection showed a smaller range among genotypes from 36.8% (Accession G41) to 92.0% (Accession G92) with a mean of 71.6%.

### Analysis of mildew tolerance with linear mixed models (LMM)

The severity of infection and sporulation measurements are expressed as proportions. We therefore fitted the LMM in Equation 1 with both the raw data and after a transformation with logit and angular functions, which are frequently used with proportions. Our goal was to assess the effect of data transformation and inclusion of check varieties on estimates of variance components, heritabilities, and genotype means The combination of these parameters resulted in six LMMs that were fitted to the traits severity and sporulation (Table 1). Genotypes were fitted as fixed effects in all models to estimate genotype means and to test for a genotype effect on disease traits. A REML ratio test showed that a heterogeneous error variance structure for the experiments provided a better model fit (*p <* 0.05) except for a single model (Table 1).

**Table 1:** Linear mixed models used to analyse quantitative response variables severity of infection and sporulation. Type of error variance refers to error variance structure between experiments. *p*-value of a REML ratio test comparing a null model with homogeneous variances of the error with a model with a heterogeneous variance structure

| Trait | Data transformation | Checks included | Variance type<br>$p$-value |
| --- | --- | --- | --- |
| Severity | No transformation | Yes | $1.72 \times 10^{-10}$ |
| | | No | $1.92 \times 10^{-04}$ |
| | Arcsine root | Yes | $4.68 \times 10^{-07}$ |
|  |  | No | 0.0014 |
| | Logit | Yes | $2.44 \times 10^{-08}$ |
| | | No | $9.77 \times 10^{-06}$ |
| Sporulation | No transformation | Yes | $1.45 \times 10^{-05}$ |
|  |  | No | 0.0390 |
|  | Arcsine root | Yes | 0.0049 |
|  |  | No | 0.199 |
|  | Logit | Yes | 0.0019 |
|  |  | No | 0.0259 |

Untransformed data for severity of infection did not strongly deviate from normality in a histogram of residuals and a QQ-plot (Supplementary Figure SS2A and B). On the other hand, a residual vs. fitted plot shows increasing variance along the *x*-axis and indicates heterogeneity of variances (Supplementary Figure SS2C). One source of variation is the experiment (Supplementary Figure S2D), which is consistent with the results of the REML ratio tests in Table 1.

A Wald test for fixed effects of genotypes on severity of infection was highly significant for all six fitted models and tests without check varieties have considerably lower *p*-values (Table 2).

**Table 2:** Wald *F*-test for the genotype fixed effects in a linear mixed model<u>an</u>alysis and mean standard error of the difference (s.e.d.), genetic variance 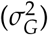 and heritability 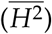 estimates from a linear mixed model for the traits severity and sporulation and a generalized linear mixed model for the trait incidence.

|  |  | Check | LMM analysis |  |  | Mean |  |  |
| --- | --- | --- | --- | --- | --- | --- | --- | --- |
| Model | Data transformation | included | F-value | d.f. | p-value | s.e.d. | $\sigma_G^2$ | $\overline{H^2}$ |
| *Trait: Severity of infection* |  |  |  |  |  |  |  |  |
| 1 | Untransformed | Yes | 3.573 | 131 | $1.18 \times 10^{-18}$ | 0.12 | 0.02 | 0.72 |
| 2 | | No | 23.95 | 130 | $3.56 \times 10^{-93}$ | 0.12 | 0.03 | 0.77 |
| 3 | Arcsine root | Yes | 3.79 | 131 | $3.35 \times 10^{-20}$ | 0.14 | 0.03 | 0.74 |
| 4 | | No | 28.95 | 130 | $8.41 \times 10^{-103}$ | 0.14 | 0.03 | 0.78 |
| 5 | Logit | Yes | 3.75 | 131 | $6.30 \times 10^{-20}$ | 0.58 | 0.46 | 0.73 |
| 6 | | No | 30.08 | 130 | $5.59 \times 10^{-104}$ | 0.59 | 0.61 | 0.77 |
| *Trait: Sporulation* |  |  |  |  |  |  |  |  |
| 1 | Untransformed | Yes | 4.47 | 131 | $4.39 \times 10^{-25}$ | 0.16 | 0.04 | 0.78 |
| 2 | | No | 8.37 | 130 | $1.55 \times 10^{-44}$ | 0.17 | 0.05 | 0.78 |
| 3 | Arcsine root | Yes | 4.89 | 131 | $7.30 \times 10^{-28}$ | 0.20 | 0.07 | 0.79 |
| 4 | | No | 11.35 | 130 | $1.13 \times 10^{-57}$ | 0.20 | 0.08 | 0.80 |
| 5 | Logit | Yes | 4.96 | 131 | $2.47 \times 10^{-28}$ | 0.87 | 1.51 | 0.80 |
| 6 | | No | 11.43 | 130 | $4.67 \times 10^{-58}$ | 0.91 | 1.85 | 0.82 |
| *Trait: Incidence* |  |  |  |  |  |  |  |  |
|  |  | Yes | 1.62 | 131 | 0.0003 | 0.55 | 0.10 | 0.4 |
|  |  | No | 2.39 | 130 | < 0.0001 | 0.48 | 0.08 | 0.4 |

Estimation of variance components allows to model sources of variation and to account for the structure of an experimental design (Harrison *et al*., 2018). For the trait severity of infection, proportions of variance components were highly similar among models. We obtained the highest estimates and confidence intervals for between experiments variance 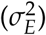 and genotype by experiment interaction variance 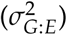 components. Estimates of variance components for experiment 1 to 3 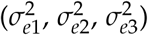 and variance of blocks nested within experiments 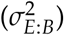 were much lower (Supplementary Figure S3 A-C, Supplementary Table S2). Estimated variance components were highly similar in a comparison of models with and without checks because the 95% confidence intervals overlapped, but variances of models with checks were smaller than without checks (Supplementary figure S3 A-C).

The results for the trait sporulation after infection are very similar to severity after infection. The Wald F-test for fixed effects was highly significant in every model fit with sporulation as response variable, which indicates that host genotypes differ in sporulation (Table 2). Removal of check varieties strongly reduced *p*-values. For sporulation, the largest variance component in every model was genotype by experiment interaction, 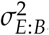, but in contrast to severity of infection, estimates of variance between experiments 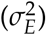and variance of blocks within experiments 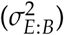 were the lowest among all variance components in all models (Supplementary Figure S3D-F). Taken together these analyses provide strong evidence for an effect of genotypes on severity of infection and sporulation that is robust with respect to the data transformation and the effects of a blocked design.

### Generalized linear mixed model (GLMM) analysis of incidence data

For the trait incidence of infection, we used a GLMM because it allows to fit non-normally distributed data like discrete proportions and to include random effects. We used a *logit* link function and assumed homogeneous variances between experiments as indicated by the conditional Pearson residuals, i.e, there is no sign that the experiments are sources of variation that need to be accounted for (Supplementary Figure S4). Using incidence as response variable in two GLMMs that differed by the inclusion and exclusion of check varieties, the test for fixed effects in both models was significant (*p <* 0.001). The genotype by experiment 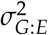 variance component was larger in the model with checks (Figure 1). Variance components reflecting the experimental design, 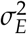 and 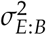, have the largest standard errors in both models. Additionally, in the GLMM without the checks the experiments 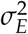 variance was zero, indicating that they are comparable between each other. The latter variance component and the residual error variance components were the largest regardless of the model used. In summary, like the other two disease traits, incidence of infection also shows a strong effect of genotypes on trait variation.

**Figure 1:**
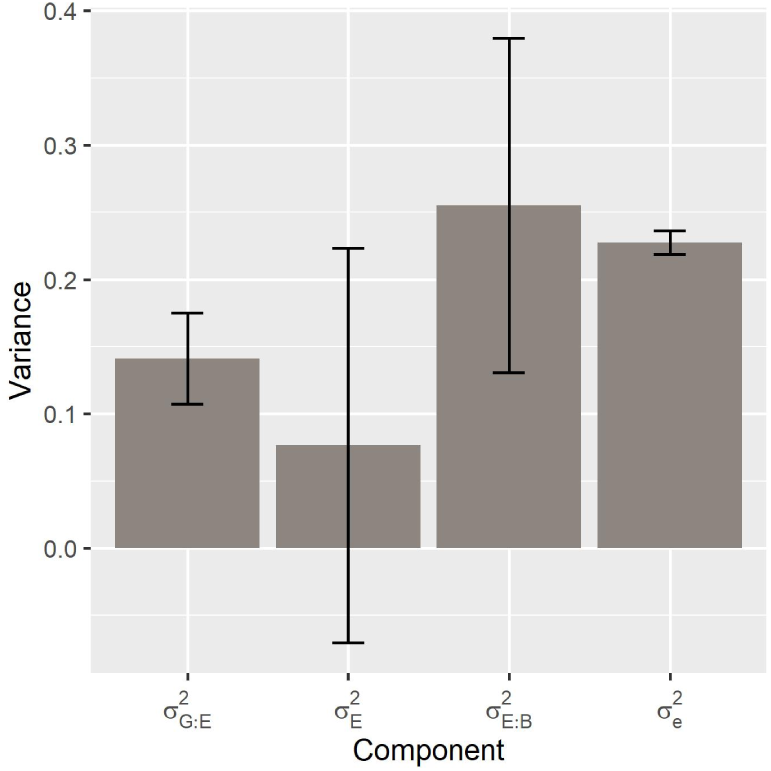
Variance component estimates and their standard errors for incidence of infection in a GLMM with check varieties. 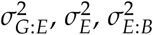: Variance components for the genotype by experiment interaction, experiments and blocks nested within the replicates, 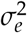 : Residual variance.

### Correlations between traits

The similar variance component structures of the three disease traits (Figure 1 and Supplementary Figure S3) suggests that they are correlated. Adjusted means of the traits severity of infection and sporulation are highly correlated with checks (*R* = 0.91, *p <* 0.001; Figure 2) and without checks included (*R* = 0.9, *p <* 0.001). The correlation of mildew incidence with both severity and sporulation was markedly lower (*R* = 0.67 and *R* = 0.65, respectively, *p <* 0.001).

**Figure 2:**
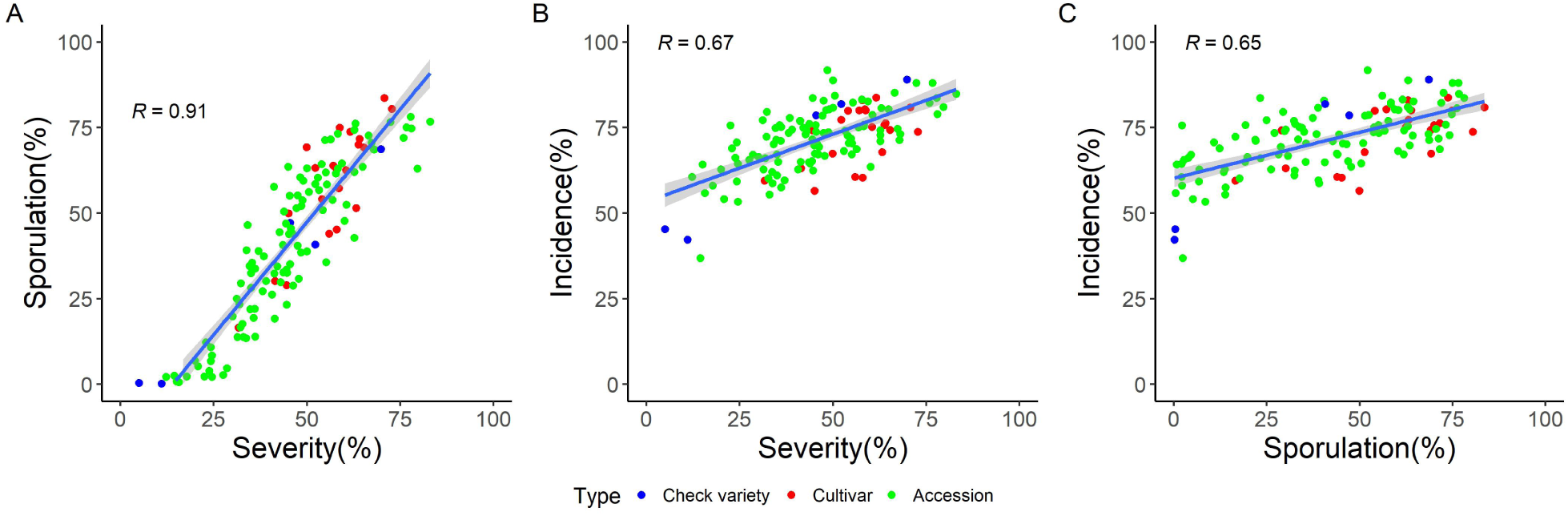
Correlations between percentage of sporulation and severity of infection (A), severity and incidence of infection (B) and sporulation and incidence of infection (C). In all three cases, the correlation was highly significant (*p <* 0.0001)

### Analysis of heritability

We also estimated heritability, 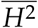 of the three disease traits and evaluated the effect of transformations on these estimates. The mean standard error of the difference (s.e.d.), which measures precision of pairwise comparisons in each model, and genetic variance, which is estimated when genotypes are fitted as random effect, are both components of 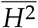 (Equation 3). Both parameters showed a small increase between models with and without checks, and we observed this difference with all data transformations (Table 2). Higher mean s.e.d values show that a removal of checks decreased the precision of pairwise comparisons. In consequence, 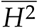 estimates in models without checks or with an arcsine root transformation resulted in marginally higher estimates than models with checks or with other transformations, respectively (Table 2). To summarize, data transformations and the exclusion of replicated checks have a little effect on the estimation of 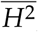, because heritability estimates remain within a narrow range from 0.72 to 0.78 for severity of infection, and a range from 0.78 to 0.82 for sporulation for all models analysed. For incidence of infection, estimated heritability was 0.40, the genetic variance was 0.10 and the mean s.e.d was 0.55 in the GLMM model with checks and 0.08 and 0.48 in the model without check varieties.

### Ranking of genotypes by mildew tolerance

The strong effect of genotype on the three pathogen traits suggests that genebank accessions and varieties in our sample are highly variable with respect to mildew tolerance. We therefore compared means in all three traits and identified substantial differences between genotypes. Adjusted mean values for severity of infection range from 5% to 83% in the LMM of untransformed data and with checks, and show a very similar range in the other models. Models without check varieties result in a smaller range for this trait because the two check varieties *C. album* and Puno had the lowest estimates for severity (Figure 3A). We did not observe a strong effect of check varieties and the type of data transformation on the ordering of genotypes for severity of infection. Therefore, differences between genotypes for this trait are robust and allow to identify tolerant and susceptible genotypes. The check varieties *C. album*, Puno and genebank accessions G41, G42, G76, G93, G96 and G112 are most tolerant, whereas check variety Vikinga, cultivars CV13 and CV21, and accessions G4, G9, G57, G67, G82 and G91 are most susceptible (Figure 3A).

**Figure 3:**
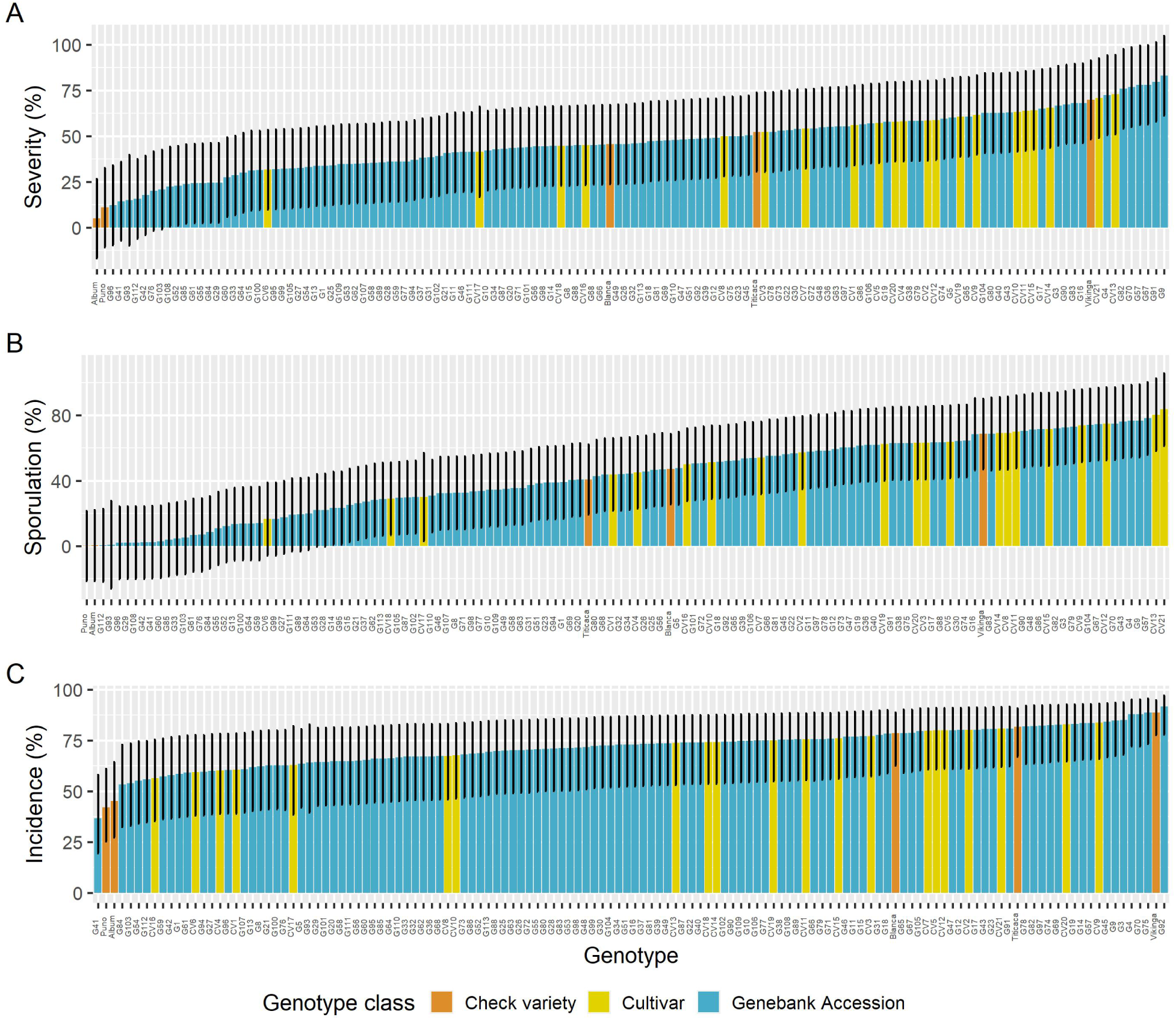
Estimated means of severity (A), sporulation (B) and incidence of infection (C) ordered from small to large for the genotypes after fitting a (G)LMM model with untransformed data (A,B) and checks included. Error bars represent 95% confidence intervals.

The genotypes show a similar pattern with sporulation. The adjusted means of untransformed sporulation estimated including the checks ranged from 0.2% (Puno) to 83.6% (CV21). Transformations of the sporulation data had small effects on the distribution of the estimated means, whether checks were included (Figure 3B). Although the arcsine root transformed data seems to have a larger range. The most tolerant genotypes for sporulation are the check varieties *C. album* and Puno, and genebank accessions G29, G41, G42, G93, G96, G106, G108, G112. The most susceptible genotypes are check variety Vikinga, cultivars CV12, CV13, and CV21, and accessions G4, G9, G43, G67, G70 and G104 (Figure 3 B).

The variation among genotypes for the trait incidence is lower than for the other two disease traits (Figure 3C). Adjusted mean values were little affected by the inclusion or exclusion of checks. Genotypes with low incidence include check varieties Puno, *C. album* and accession G41; while check variety Vikinga and genotypes G75 and G92 had the highest incidence percentages (Figure 3 C).

### Relationship of disease traits with genebank passport data and seed saponin content

The genotypes accessions included in the experiment were selected using information on mildew tolerance from passport data and INIAF to obtain a set of accessions, which is polymorphic for this trait. The severity and incidence data recorded in the passport data of genebank accessions are highly incongruous with our results. For example, 35 of 106 accessions were recorded under the 0% severity category in the passport data but no accession was classified as such in our analysis; 16 accessions were assigned to 0.1-25% group and 14 in this study; 26 accessions as 26-50% vs. 56 in our dataset. According to the passport data, 26 accessions are in the most susceptible category (75-100%), and only 6 accessions in this study.

The only significant correlation of disease and stomata traits was between severity and stomata width (*r* = 0.18, *p* = 0.041). We also tested whether saponin content in seeds is correlated with disease susceptibility and carried out foam tests with seed harvested in two locations because saponin content varies between phenological stages and environments. 105 genotypes had seed available from both locations, Bolivia and Denmark, while 26 genotypes were harvested only in Denmark. Foam height measurements were not correlated between sources of seed (Pearson’s *r* = 0.16, *p* = 0.11). In a comparison of estimated average severity and sporulation of genotypes with and without saponins, we did not observe any systematic pattern based on the seed source or if accessions or cultivars were compared separately (*t*-test with *p >* 0.05).

### Isolation from *C. album* and cross-infection on *C. quinoa*

The isolate of *P. variabilis* used in this study was originally isolated from *C. album* and afterwards vegetatively propagated on *C. quinoa* (cv. Blanca and Vikinga). Spores harvested from these plants and inoculated onto *Chenopodium* spp. showed low disease severity (4%) and lowest sporulation (0.4%) to *C. album* compared to the *C. quinoa* genotypes (Figure 3). This is the first time that cross infection from *C. album* has been reported. The BLAST comparison of the ITS DNA sequences used to validate the isolate showed a 100% match to an isolate obtained from *C. quinoa* cv. Atlas collected in 2001 (EU 113305).

### Whole genome sequencing of accessions

Given the highly significant genotypic effect on response to mildew infections, we sequenced a subset of 88 accessions and cultivars representing the range of phenotypic variation to conduct a GWAS. The sequence genotypes included two control varieties (Puno and Titicaca), 18 cultivars, and 68 genebank accessions of which 39 originated from Bolivia, 19 from Peru, two from Ecuador and 1 from Chile. The average severity of the sequenced genotypes ranged between 11% and 83%, with an average of 47%.

Sequencing produced 7.9 *×* 10^11^ bp in 2.6 *×* 10^9^ reads. After mapping processed sequence reads to the quinoa reference genome version 1, sequence coverage per genotype ranged from 0.38X (Check variety Titicaca) to 9.17X (Accession G37) with an average of 3.24X. The proportion of mapped reads per sample ranged from 99.3% to 99.9%, with an average of 99.8%. In contrast to the nuclear genome, the chloroplast and mitochondrial genomes were overrepresented in our sample, with a coverage of 109X and 32.6X, respectively.

Variant calling with GATK identified 18, 017, 831 biallelic SNPs across the genome. After filtering for minor allele frequency and sample missingness, and testing for departure from Hardy-Weinberg Equilibrium 4, 131, 562 variants remained. We imputed genotypes with coverage *<* 8X, and sample and position missingness below *<* 0.7, which produced 606, 791 SNPs from 61 samples with an estimated accuracy of 96.9% based on 5, 000 hidden markers. Figure 4A shows that linkage disequilibrium, measured as *r*^2^, drops to 0.1 within 22-25Kb in the sequenced genotypes.

**Figure 4:**
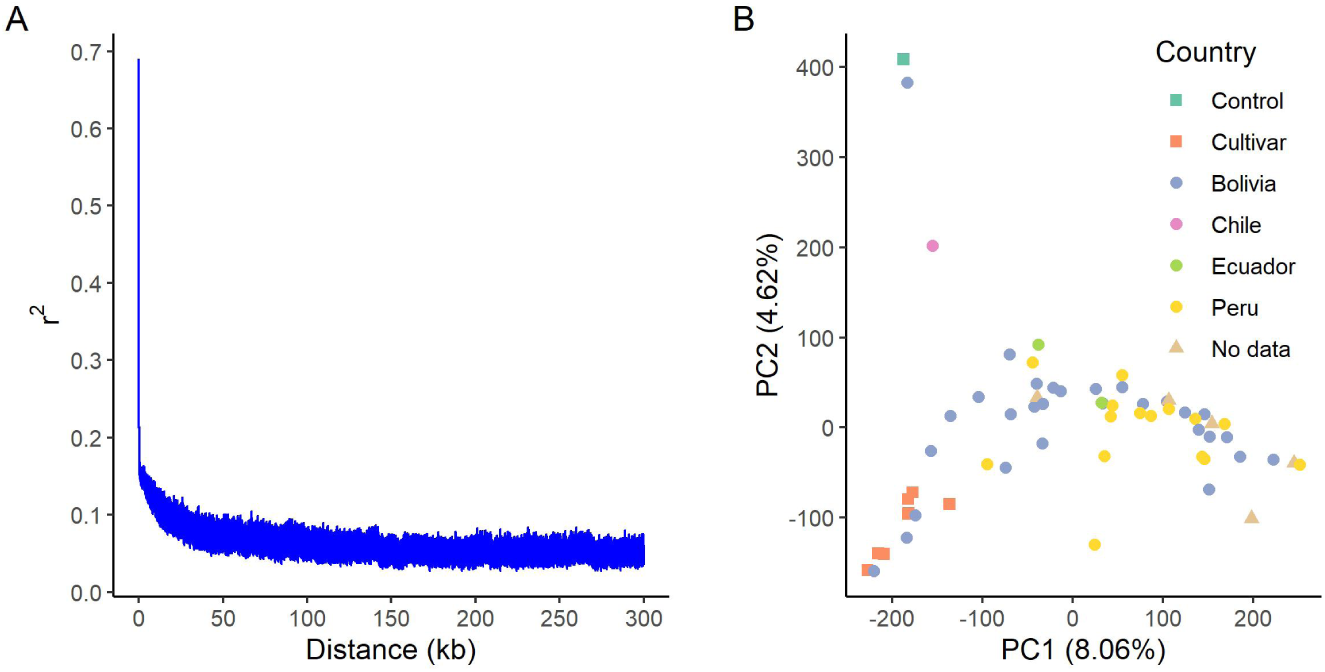
(A) Decay of linkage disequilibrium (LD) expressed as a function of physical distance (kb) and *r*^2^. (B) Plot of the first two principal components of a PCA analysis. Percentages in each label are the proportion of variance explained by each PC; colors indicate the country of origin as indicated in the passport data.

The principal component analysis (PCA) shows that all genotypes classified as cultivars cluster together (Figure 4B, lower left corner). The check variety Puno and the genebank accession G71 (originating from Bolivia) are separated from the other genotypes. Accession G101 is also separated from the major group and originates from Chile, suggesting it is of the coastal ecotype. Accession G42, which appears to be separate form the main group (Figure 4B, middle bottom), has very low severity and sporulation, which is comparable to the variety Puno. The remaining accessions originating from Bolivia, Peru and Ecuador are mixed and do not form distinct groups.

### Association mapping for severity of downy mildew

We carried out two different GWAS analyses to detect genomic regions associated with severity of downy mildew infection. Since severity and sporulation showed a high degree of correlation, we conducted the GWAS only with the first of the two traits. An analysis of 603, 871 SNPs in 61 genotypes with FarmCPU did not uncover statistically significant associations with average severity when the model was fit with or without principal components (PCs) (Figure 5A and Supplementary Figure SS5A, respectively). A single variant on chromosome 4 (S04 33782670) is located above a threshold (1.656 *×* 10^*−*06^) in the model without correction for population structure using principal components (Figure 5). The QQ plots showed no sign of inflation or deflation of *p*-values with respect to the theoretical expectation (Figures 5B and Supplementary figure SS5B, respectively) and therefore supports the absence of a significant association.

**Figure 5:**
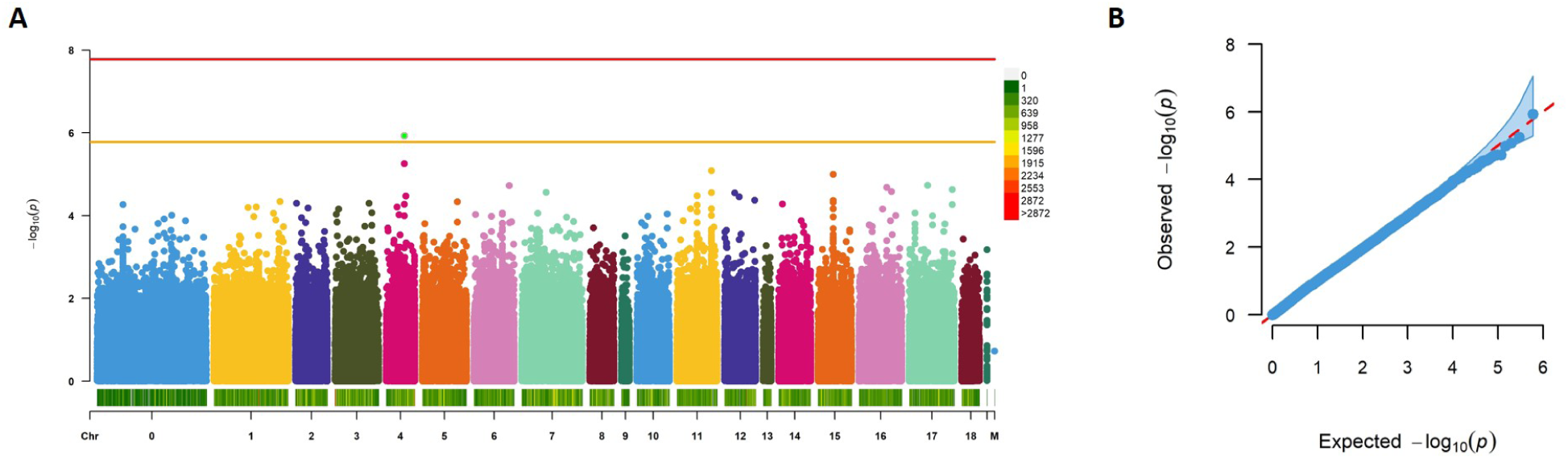
Association mapping for downy mildew severity using FarmCPU without principal components as covariates. Manhattan plot (A). Red line shows the Bonferroni corrected threshold for *p* = 0.01 and orange line indicates a suggestive threshold (1/number of markers). Bar at the bottom indicates marker density. QQ plot (B) for the FarmCPU model with the 95% confidence interval (light blue); Red line draws the expected distribution of *p*-values.

We also used a *k*-mer based approach because it allows the inclusion of additional genotypes with lower sequence coverage (*n* = 88) and is not biased to genomic regions included in a reference sequence. This analysis was based on an average of 570, 741, 731 *k*-mers of length 31 per sample. The check variety Titicaca (67, 365, 628) and genebank accessions G37 (814, 316, 239) were the genotypes with the lowest and highest numbers of *k*-mers counts, respectively. In total, 992, 946.265 *k*-mers passed the filters and used for both kinship matrix estimation and the subsequent GWAS. For the first stage of the GWAS, 880, 137, 481 *k*-mers were tested and 10, 001 passed the first filter to be fit with GEMMA. The smallest *p*-value for any *k*-mer was 9.19 *×* 10^*−*10^ for a single *k*-mer. Therefore, this analysis also did not detect any significant associations with the trait severity of infection given a permutation-based 5% *p*-value threshold of 1.505 *×* 10^*−*10^ for the *k*-mer analysis.

## Discussion

The inoculation of quinoa varieties and genebank accessions with an isolate of the downy mildew pathogen *Peronospora variabilis* revealed substantial variation in the three infection related traits severity of infection, sporulation and incidence of mildew (Figure 3). Using a mixed model approach we validated that estimates of genetic effects, variance components and heritabilities are robust with respect to data transformation and the inclusion or exclusion of check varieties between experiments. The differences in susceptibility to mildew infection have a strong genetic component as indicated by high genetic variance component estimates and high heritabilities, but we were not able to identify individual genomic regions strongly associated with mildew susceptibility in a GWAS. In previous studies, ecotype, environmental or physiological parameters like altitude of site of origin, seed saponin content, or size and density of stomata were postulated to be correlated with disease tolerance. We did not find any strong correlation of the three disease traits with any of these parameters except for width of stomata.

### High genetic effect of mildew tolerance

Mildew tolerance was scored in a single plant per block, in the case of accessions and cultivars, while check varieties had multiple plants because one person scored each leave of all plants and the labor-intensive process has a time limitation for scoring window (max. 12 hrs). To evaluate the robustness of parameter estimates, we used linear mixed models and various combinations of data transformation and inclusion or exclusion of check varieties. Although the use of data transformation is under debate (Milligan, 1987; O’Hara and Kotze, 2010; Warton and Hui, 2011), it did not have large effect on the distribution of residuals or on the tests of fixed effects. This provides strong evidence for the robustness of our estimates because we phenotyped approximately the same number of plants per genotype, which reduces the effect of heteroscedasticity (Zimmerman, 2006), and from the robustness of LMMs to heterogeneous error variances (Jacqmin-Gadda *et al*., 2007). The effect of replicated checks in the three experiments on model fit and parameter estimation was minor, because their removal caused only a small increase in 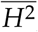 estimates. This robustness results from a balanced experimental design, in which changes to differences are only small (Piepho and Möhring, 2007).

A limitation in fitting LMM and GLMM is that reliable estimation of the variance of a random effect requires at least five levels, i.e. locations, experiments, years, etc. (Harrison *et al*., 2018). The low number of groups in our experiments explains the large confidence intervals of the design variance components 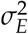 and 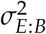 (Supplementary tables S2 and S3), which therefore are not reliable estimates of variation between experiments or blocks. We nevertheless modeled these effects as random to use inter-block information and to account for non-independence of the data, because genotypes were nested in blocks, which were nested within experiments (Bolker *et al*., 2009; Harrison *et al*., 2018).

Previous work on quinoa yield traits found high estimates of genotype-by-environment variance components (Bertero *et al*., 2004). The genotype-by-experiment interaction variance 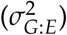 in our study was also large in comparison to other variance components across traits and models, which reflects that even subtle differences in the environment can cause a genotype to respond differently to the disease. From a plant breeding perspective, the 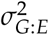 variance component is important because it masks the genotypic component of phenotypic variance. A high observed 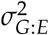 suggests that future studies of mildew tolerance using diverse quinoa genotypes should include multiple locations and years (Hayward *et al*., 1993), or unbalanced designs with replicated checks that allow testing of many genotypes (Singh and Bhatia, 2017).

### High correlations between disease-related traits

To identify different reactions of host plants, we measured three different disease-related traits. A comparison of severity of infection and sporulation is of interest because pathogen populations typically harbor high levels of genetic variation for both virulence and fecundity (Sacristán and García-arenal, 2008; Barrett *et al*., 2009). Some quinoa host genotypes might allow fast fructification of the pathogen while others may suppress its proliferation (Mhada *et al*., 2015). We found a very strong positive correlation between severity and sporulation and scoring of the former seems to be a good predictor of the latter. For this reason severity alone can be used to assess a panel of genotypes because it also showed a smaller error variance than either sporulation (Figure S3D-F) or incidence (Figure 1). The correlation coefficients of the disease traits in our study (*R* = 0.9) is larger than the coefficient reported in an evaluation of scoring methods for downy mildew in cucumber (Pitrat, 2008). However, sporulation should be measured in highly resistant genotypes with *<* 5% severity of infection, because in such genetic backgrounds pathogen proliferation may be strongly impaired as observed in the wild relative *C. album*, check variety Puno and several genebank accessions.

### Analysis of heritability

Consistent with estimates of genetic variance, heritability estimates were moderately high and very similar between models with a range from 0.72 to 0.78 for severity and from 0.78 to 0.81 for sporulation. In comparison, estimated heritability of sporulation of downy mildew in grapevine (*Vitis vinifera*) was around 0.40 (Divilov *et al*., 2018), and resistance to systemic infection by sorghum downy mildew in maize was in the range of 0.61-0.68 (Lohithaswa *et al*., 2015). Although estimates of 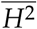 from greenhouse experiments may differ from field trials (Gardner and Latta, 2008), the high heritabilities for the disease traits indicate the selection for higher mildew tolerance is possible. In this respect our results are consistent with previous estimates of 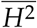 for physiological, morphological and yield traits, which are also high (*≈* 0.85) (Santis *et al*., 2016) and indicate that multiple traits of quinoa can be substantially improved by plant breeding.

### Identification of accessions tolerant to downy mildew

Although our panel includes the main quinoa ecotypes (Tapia *et al*., 1979), there was no correlation between elevation and severity of infection or sporulation. Since infection requires high humidity (Figure 8), a lack of such a correlation reflects microclimatic variation of humidity in high altitudes. For example the Bolivian highland (Altiplano) is more humid in the North than in the South (Danielsen *et al*., 2003). In addition, ecotypes from the Andean Valley (2,500-3,500 masl), where humidity also tends to be higher, are among the most tolerant accessions. (Danielsen and Ames, 2000; Danielsen *et al*., 2003; Gabriel *et al*., 2012). These observations are consistent with our results because five accessions from the southern altiplano had the highest severity of infection (G4, G9, and G82 from Bolivia, and G67, and G99 from Peru).

We observed that a ranking of genotypes by their mildew tolerance is robust with respect to disease traits and analysis methods, which allows to identify accessions high tolerance for further investigations or utilization in breeding. Tolerant genotypes include the wild relative *C. album* and the Puno check variety, which was one of the genotypes with the lowest severity, sporulation, and incidence of downy mildew (Figure 3A-C), as well as a set of Bolivian cultivars and genebank accessions. For example, cultivars Mañiqueña (CV21) and Phisankalla (CV10) perform well in dry areas like the southern Altiplano in Bolivia, but are susceptible to mildew in humid environments (Danielsen and Ames, 2000; Gandarillas *et al*., 2015a; Murphy *et al*., 2018). This phenotype was confirmed in our study because both cultivars are among the most susceptible under the humid conditions of our experiment (Figure 3). It has been proposed that genotypes with good performance in dry and a high disease susceptibility in humid environments either have not been selected for disease tolerance during domestication or have an advantage in dry environments possibly because of the cost of resistance (Danielsen *et al*., 2003). Current evidence is contradictory and does not establish such a relationship, because cultivar ‘Rosa Blanca’ (CV6) was developed in a dry region (Gandarillas *et al*., 2015a), but has a higher tolerance to the disease with an average severity of 32%, whereas the more susceptible cultivars ‘Jach’a Grano’ (CV15; 64% severity) and ‘Aynoka’ (CV20; 58% severity) originated from the same breeding program. Such a high variation may result from the interaction between genotypes and experiments (Figure S3) and is consistent with similar GxE interactions anatomical and yield-related traits of *C. quinoa* (Santis *et al*., 2016; Al-Naggar *et al*., 2017).

An important limitation of our study is the use of a single isolate only of *P. variabilis* for inoculation because it does not allow to test whether mildew tolerance is race-specific or reflects a quantitative resistance. Preliminary evidence supports the latter hypothesis, because cultivars Kurmi (CV16) and Mañiqueña Real (CV21) were inoculated with a Bolivian isolate of *P. variabilis* and classified as tolerant and susceptible, respectively, as evaluated by the disease progression of downy mildew *P. variabilis* (Rollano-Peñaloza *et al*., 2019). Our results confirm the differences between these two cultivars with a different isolate because Kurmi (45.1% severity of infection) was less susceptible than Maniqueña (70.7%) (Figure 3). Since Kurmi was developed for cultivation in the highlands (3,600-3,800 masl) and selected in field trials for downy mildew resistance (Bonifacio, 2015), a high tolerance of Kurmi to two different isolates supports a quantitative disease tolerance. Our results also support previous work that found recombinant inbred lines from both Chilean and Peruvian origin segregating for quantitative mildew resistance in a F_2:6_ population (Benlhabib *et al*., 2016).

Comparison of passport data with our results is limited by missing information on scoring methods, phenotypical stage of the plant, genetic constitution of accessions (e.g., extent of heterozygosity) and information about field trials in Bolivia. In addition, our data on severity of infection is based on one pathogenic isolate whereas the passport data is based on natural infections of local races whose virulence might differ from the Danish isolate used in this study (Danielsen and Lübeck, 2010). Previous studies have shown that Andean isolates are genetically distinct when highly sensitive polymorphism methods of identification are used (Danielsen and Lübeck, 2010). Based on the PCR sequencing data, our isolate showed complete sequence identity in the ITS region to specimen EU 113305 collected in Tåstrup, Denmark on C. quinoa cv. Atlas in 2001 (Choi *et al*., 2010).

### Relationship of disease traits with stomatal traits and seed saponin content

*P. variabilis* enters host tissues through stomata, and its haustoria emerge from the stomatal pore to release spores (Kitz, 2008; Choi *et al*., 2010), which may explain the preference of pathogen for humid conditions because stomata are typically open under such conditions (Lange *et al*., 1971). We therefore tested whether pathogen traits are correlated with stomatal traits and found that only severity of infection showed a weak correlation with stomatal width. This result should be taken with caution because our measurements were based on only a single leaf per genotype due to the high effort required to obtain the data. Future phenotyping should make use of automated image analysis methods to obtain larger data sets for stomatal traits.

Downy mildew remains dormant in the pericarp of the seed (Danielsen *et al*., 2004) and the saponin content of the seeds could influence the response of a genotype during the early stages of infection. However, we found no support for a relationship between saponin content of the seed and mildew severity. One explanation for the absence of a correlation may be that the foam test revealed GxE effects, because the scores for content of saponin differed between seed samples of the same genotypes obtained from plants cultivated in different locations. This is expected because the content of saponin is variable over time and depends on the water status of the plant (Solíz-Guerrero *et al*., 2002; Martínez *et al*., 2009).

Therefore, our results do not support the hypothesis that mildew tolerance is substantially influenced by other traits such as stomata characteristics or saponin content.

### Association mapping for severity of downy mildew

We conducted a GWAS with severity of mildew infection and whole genome resequencing data to test whether the observed differences between genotypes are caused by few genomic regions. Both GWAS methods sed failed to detect significant associations of variants or *k*-mers with severity of infection. The power of GWAS depends on the sample size of the association panel and on the genetic architecture of a trait of interest (Korte and Farlow, 2013). Our analysis was limited by a small sample size of 61 (FarmCPU) and 88 (*k*-mer analysis) accessions, and possibly by the genetic architecture of severity of infection because distribution of phenotypic values suggests it is a polygenic quantitative trait (Figure 3A). A polygenic response of quinoa to *P. variabilis* infections is supported by multiple studies that include greenhouse experiments and field trials (Gandarillas *et al*., 2015b; Benlhabib *et al*., 2016; Kumar *et al*., 2006; McElhinny *et al*., 2003; Mhada *et al*., 2015; Curti *et al*., 2014; Khalifa and Thabet, 2018).

Our results provide a perspective for a more efficient resistance breeding in quinoa. The large variation found in mildew tolerance and high heritabilities of disease traits allows the development of QTL mapping populations by crossing genotypes from both ends of the distribution (e.g., Danish varieties Puno and Titicaca; Figure 3). Previous work suggests that QTL mapping may identify major R genes that could be useful in quinoa breeding because mildew tolerance is modulated by incomplete gene effects (Poland *et al*., 2009), which depends on pathogen agressivity, as observed on Ecuadorian material (Ochoa *et al*., 1999). Furthermore, the segregation ratio of mildew severity in an *F*_2_ mapping population derived from a cross of bitter and sweet (i.e., no seed saponins) genotypes suggested that mildew tolerance shows a dominant inheritance (Mastebroek and van Loo, 2000).

The analysis and utilization of genetic variation for mildew tolerance will be enhanced by more high throughput phenotyping methods. Our experimental setup was on targeted inoculations with a single isolate, which contributes to a robust and repeatable estimation of disease tolerance, but it is work intensive and limits the number of genotypes for genetic mapping. However, alternative approaches such as scorings of detached leaves or randomized selection of leaves on the field can be misleading because host genotype x pathogen genotype x environment (GxGxE) effects in the field are difficult to control. Furthermore, symptoms of pathogen infection are influenced by the position and age of leaf tissue (Calixtro *et al*., 2017) which results from induced resistance that occurs not only at the site of the initial infection but also in distal, uninfected parts (Conrath, 2011). Therefore, in addition to a controlled greenhouse experiment used in this study, multilocation field trials of segregating populations that use modern phenotyping technologies such as deep learning to score pathogen infections are a complementary approach in resistance breeding (Sperschneider, 2019). Both approaches in combination with genetic analysis will contribute to the development of improved quinoa varieties in both within and outside of the native cultivation range.

## Conclusion

Our study revealed a high level of variation of quinoa varieties and accessions to *Peronospora variabilis* infections. We have shown that cross-infection from *C. album* to *C. quinoa* and vice-versa is feasible and this widely distributed weed is an alternate host for the *P. variabilis* with implications for quinoa cultivation. The substantial variation in mildew tolerance between genotypes has a strong genetic component is therefore is amenable to selection in breeding programs. However, inferences based on a single experiment - or a single location field trial - should be taken with care because a large genotype by experiment interaction was found, so future work on the resistance of *C. quinoa* to *P. variabilis* must take this into consideration during the design and planning phases.

## Methods

### Plant material

The quinoa genotypes analyzed in this study consist of 106 accessions stored in the National Germplasm Bank of Bolivia. They include landraces collected in Bolivia (55 accessions), Peru (33), Ecuador (7) and Chile (4), in altitudes ranging from 2 masl to 4,082 masl (Figure 6). Seven accessions had no information about their origin. We also included 21 Bolivian cultivars, the Bolivian variety ‘Blanca’ (‘Blanquita’) and three Danish varieties ‘Puno’, ‘Vikinga’ and ‘Titicaca’. The list of accessions and their passport data are provided in Supplementary File 1 (Source: http://germoplasma.iniaf.gob.bo).

**Figure 6:**
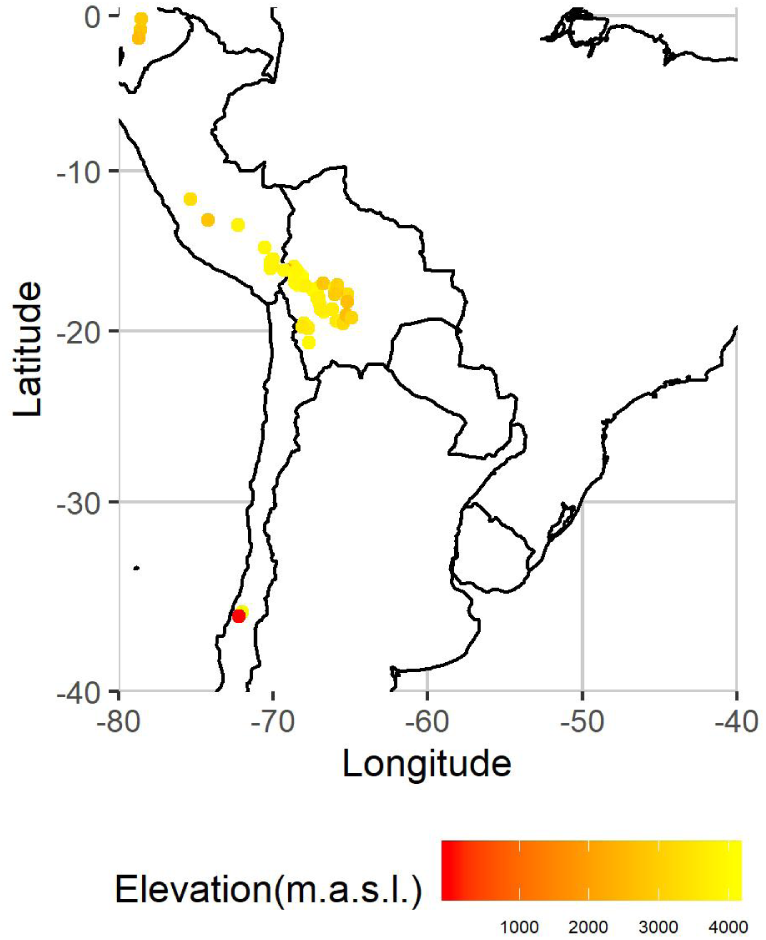
Distribution of germplasm bank accessions across south America by elevation according to the passport data. Source: Bolivian National Germplasm Bank (Source: http://germoplasma.iniaf.gob.bo)

Genebank accessions were selected to represent both the geographic diversity of quinoa and variation in mildew tolerance, which was scored in field trials in La Paz (Bolivia) based on spontaneous infections by *P. variabilis*. Additional information on the genetic status of accessions, scoring method, phenological stage during scoring, and trial locations were not available from passport data.

Four quinoa varieties and the wild relative common goosefoot *Chenopodium album L*.) were used as checks in greenhouse experiments. The four quinoa varieties include the cultivar ‘Blanca’, which is adapted to the Northern highlands and Inter-Andean valleys of Bolivia and partially resistant to downy mildew (Bonifacio, 2015; Gandarillas *et al*., 2015a). The other three check varieties ‘Titicaca’, ‘Puno’ and ‘Vikinga’ were developed in the quinoa breeding program of the University of Copenhagen. Varieties Puno (KVL 37) and Titicaca were bred from Chilean and Peruvian landraces and selected for earliness and adaptation to European conditions (Stikic *et al*., 2012; Sun *et al*., 2014). They showed different levels of downy mildew susceptibility in a field trial (S.-E. Jacobsen, personal communication), which was confirmed in a pilot experiment for this study (Supplementary Figure SS1). *Chenopodium album* is closely related to quinoa and a widely distributed weed. *C. album* seeds used in this study were collected in 2017 and 2018 at a former quinoa breeding field at the experimental station of the Faculty of Science, University of Copenhagen (Højibakkegaard, Tåstrup).

### *Peronospora variabilis* isolate used for inoculation

Previous research recognized the role of alternate hosts on the evolution and spread of pathotypes (Lewis *et al*., 2018) *P*.*variabilis* is a pathogen of both *C. quinoa* and *C*.*album* (Aragón and Gutiérrez, 1992; Choi *et al*., 2010; Danielsen and Lübeck, 2010; Testen *et al*., 2014; Kara *et al*., 2020). XXX To obtain a defined isolate of *P. variabilis*, leaves from *C*.*album* with typical downy mildew sporulation were collected in late September 2018 at a former quinoa breeding field on the research station Højibakkegaard. The isolate was inoculated for maintenance and propagation into two quinoa cultivars (Blanca and Vikinga) using a protocol by Danielsen and Ames (2000). We used these two cultivars because they differed in their latent period (Mhada *et al*., 2015). The latent period lasted five days in the Vikinga variety and 7-10 days in the Blanca variety. These differences allowed to maintain the pathogen on Blanca, and a quick propagation on the Vikinga variety.

We used DNA sequencing of the Internal Transcribed Spacer (ITS) region to confirm that the isolate was *P. variabilis*. A spore suspension (1 *×* 10^11^ spores/ml) produced from the maintained inoculum was filtered with a nylon filter with 20*µ*m pore size (Merck Millipore Ltd.) to capture *P. variabilis* spores, which were transferred to 1.5 ml microcentrifuge tubes containing glass beads (425 – 600 *µ*m) and kept on ice. 200*µ*l lysis buffer (DNAeasy Plant Mini Kit, Qiagen) were added and mycelia were pulverized with a sterile pestle. Further 200*µ*l of lysis-buffer with 4*µ*l of RNase were added. DNA was extracted with a DNeasy Plant Mini Kit (Qiagen) following manufacturer’s instructions. Primers designed to amplify a 1,150 bp fragment covering ITS-1 and ITS-2 region in members of the oomycete family Peronosporaceae, including species of Peronospora, Pythium, and Phytophthora, were amplified from genomic DNA by polymerase chain reaction (PCR) using Oomyc Fw-1: 5’ cggaaggatcattaccacac and Oomyc-Rv1: 5’ cgcttattgatatgcttaagttca as forward and reverse primers, respectively. PCR amplification was carried out with one cycle of 95°C for 3 min; 35 cycles of 94°C for 30 s, 55°C for 30 s and 72°C for 40 s, and one cycle of 72°C for 3 min. Amplification products were purified using QIAquick PCR purification columns (Qiagen) and the DNA concentrations were determined on a NanoDrop Lite Spectrophotometer. DNA sequencing of the PCR amplified ITS was performed at Eurofins Genomics. DNA sequences were submitted to NCBI (accession MT895880) and compared against the NCBI nr database using BLASTN (https://blast.ncbi.nlm.nih.gov).

### Identification of *P. variabilis* by microscopy and histopathology

The identity of the pathogen was also confirmed by visual analysis with microscopic and histopathology slides. Microscopic analysis was carried out with a sample of the *Peronospora* solution mounted on a glass slide. Histopathological slides were prepared from Blanca leaf pieces of *±*5 cm^2^ collected 7 days after infection. Leaf samples were coated with nail polish on the abaxial side and dried for 24 hours. The imprints were then removed and stained with Lactophenol Aniline Blue (Silva *et al*., 1985). Both microscopic and histopathological preparations were mounted on a Leica microscope MZ12.5 and photographed with a LEICA DFC420 camera under 40X magnification (Figure 7A,B). Additionally, live infections were captured with a digital microscope (Dino-Lite, model AM4113/AD4113 Dino-Lite, Naarden, Holland) (Figure 7C). After verification and calibration of the pathogen, the isolate was constantly propagated *in planta* on Vikinga and Blanca. The detailed protocol for the isolation and propagation of the pathogen is available in Supplementary File 2.

**Figure 7:**
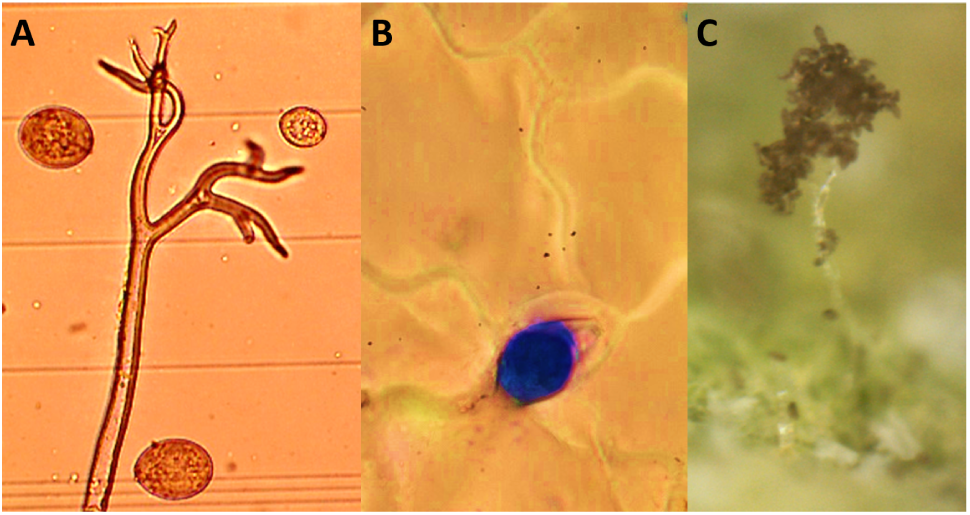
Microscopic and histopathological imaging for identificacion of *Peronospora variabilis*. A) Sporangiophore and spores of *P. variabilis* under 40X magnification. B) Spore appresorium and hyphae penetrating a stoma during infection in Blanca (7 days after infection, 40X magnification).

**Figure 8:**
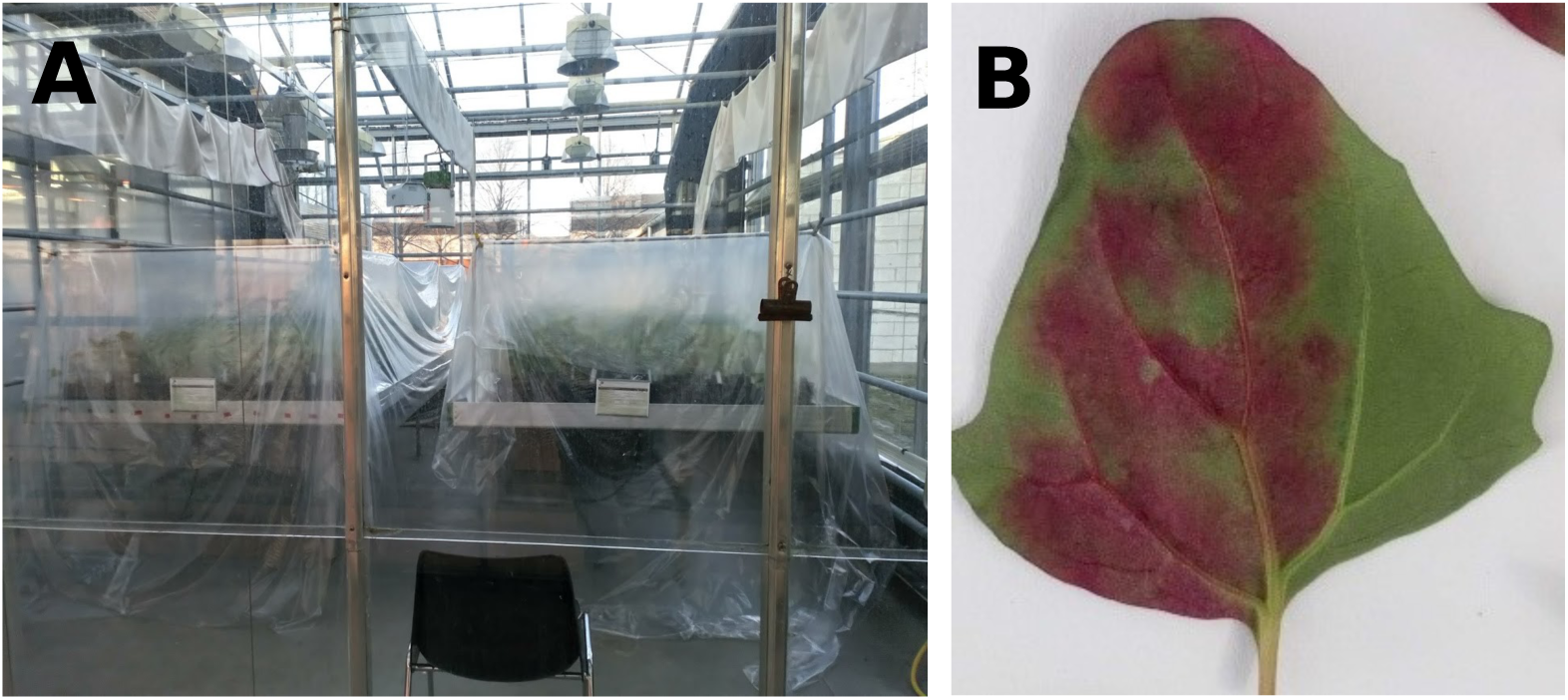
A) Two blocks of the experiment during the infection phase with plastic sheet covers to provide high humidity. B) Quinoa leaf with signs of infection by *P. variabilis*.

### Design of phenotypic characterization

Phenotypic data were collected in greenhouses of the University of Copenhagen between February and May 2019. The response of quinoa genotypes to downy mildew inoculation was evaluated in three sequential identical experiments that each included the complete set of 132 genotypes and the four check varieties with a randomized complete block design with four blocks each. Each experiment occupied a greenhouse allocated exclusively for the experiment to avoid infestations with insects or risk of cross contamination as well as the provision of biological control agents. Experiments started two weeks after the end of the prior one. Within blocks, accessions and cultivars were represented by a single plant while check varieties were represented by 2 to 5 plants.

Prior to the experiment, the Bolivian gene bank accessions were self-pollinated once to increase homozygosity because the heterozygosity of genebank accessions was unknown. Seeds were produced in a greenhouse between February and August 2017 with a 12 h photoperiod, an average temperature of 24°C during the day, 18°C at night, and irrigated with fertilized water (NPK 14-3-23 + mg EC 1.9). After the day length surpassed the plants requirements, greenhouse curtains were used to maintain the photoperiod. Seeds were harvested, cleaned and store at natural conditions until January 2018. Harvested seeds were sown in jiffy pots containing peat to assure the provision of plantlets for transplantation and grown for seven to ten days. To avoid infestation with flies, the compost was watered with a solution of gnatrol (10% v/v). Plantlets were then transplanted to 550 cm^3^ pots and grown for three weeks in greenhouses under the same conditions. The greenhouse management included biological control agents against common greenhouse-borne pests.

Three weeks after transplanting, plants were moved to different greenhouses and inoculated with a calibrated solution (1 *×* 10^5^ spores/ml) of *P. variabilis* spores and Tween 20 (1%). The solution was sprayed comprehensively onto each plant with a pressure paint gun. 50 ml of solution were used for each block. Blocks with inoculated plants were covered with a plastic sheet 5 days after inoculation (Figure8A) for 24 hours under complete darkness and a night-time temperature of 15°C to create conditions that stimulate infection. After removal of the cover, plants were grown under greenhouse conditions. Once symptoms were observed (Figure 8B), usually between 5-6 days after inoculation, plants were covered again for 24 hours to promote sporulation.

The plant response to pathogen infection was measured with the three variables: severity, sporulation, and incidence. Scoring of severity and sporulation by visual analysis of the foliar area covered by lesions of chlorotic or other color on the adaxial leaf side and the area of diseased tissue with visible spores on the abaxial leaf side, respectively. Measurements were recorded as percentages for each leaf and then averaged per plant. Incidence was calculated as the proportion of leaves with symptoms. These parameters were measured by the same person to avoid operator bias and all plants had the same age when scored.

### Phenotyping of stomata

To measure width, length, and density of stomata on leaf surfaces, a resin cast of the abaxial surface of one leaf from each genotype was made (Scarpeci *et al*., 2017). After drying, a layer of nail polish was added to the cast, left to dry, removed, mounted onto a glass slide and covered with a glass coverslip. Slides were mounted on a Leica MZ12.5 optical microscope and three fields were photographed using a Leica DFC420 digital microscope camera with 40X magnification. Width and length were measured using the Leica Application Suite software. Stomatal density was estimated as the number of stomata per unit of area and stomatal counts were obtained by using Stomata Counter, a web-based application, followed by manual curation of the data (Fetter *et al*., 2019).

### Analysis of phenotypic data with linear mixed models (LMM)

The following mixed model was used to estimate the mean severity and sporulation of the disease for each genotype in the panel, using ASREML-R package version 3.0 (Butler *et al*., 2009):

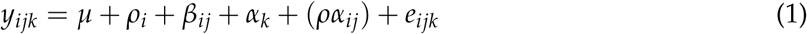

where *y*_*ijk*_ is the response (severity or sporulation) of the *k*-th genotype in the *j*-th block of the *i*-th experiment, *µ* is the general mean, *ρ*_*i*_ is the effect of the *i*-th experiment, *β*_*ij*_ is the effect of the *j*-th block nested within the *i*-th experiment, *α*_*k*_ is the genotype effect, *ρα*_*ij*_ is the genotype-experiment interaction, and *e*_*ijk*_ is the residual error term. The effects for experiments, blocks within experiments and the genotype-experiment interaction were treated as random effects because experiments were considered as a random factor and hence all effects involving a random factor need to be modelled as random (Piepho *et al*., 2003), whereas main effects of genotypes were treated as fixed. To avoid the influence of outliers on estimates of genetic variance, outliers were detected after fitting the model and removed from the dataset using the default method of the PLABSTAT package (Utz, 2001), as described in Bernal-Vasquez *et al*. (2016).

The residual error for the experiments was initially modeled as normally distributed and independent with a common variance component, where *n* is the total number of observations. In addition, we fitted a model such that the variance-covariance matrix of the vector of errors (sorted by experiments) was 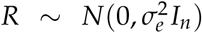, where *n* is the total number of observations. In addition, we fitted a model with independent variance components for each experiment (Isik *et al*., 2017):

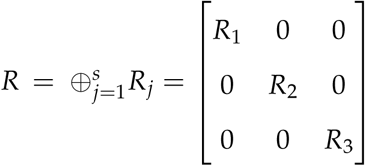

where 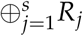 is the direct sum of matrices and *R*_1_, *R*_2_ and *R*_3_ are variance-covariance structures for each experiment, each taking the form 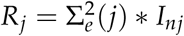, where *nj* is the number of observations in the *j−*th experiment.

### Analysis with Generalized Linear Mixed Models (GLMM)

We used a Generalized Linear Mixed Model (GLMM) to analyze the incidence of downy mildew in quinoa and fit the following model with the PROC GLIMMIX procedure of SAS software:

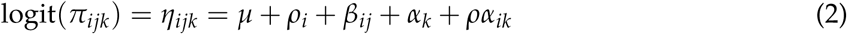

where logit is the link function between the linear predictor and the observations (*p*_*ijk*_), *ρ*_*i*_ is the effect of the *i*-th experiment, *β*_*ij*_ is the effect of the *j*-th block nested within the *i*-th experiment, *α*_*k*_ is the genotype effect, and *ρα*_*ik*_ is the genotype-experiment interaction. The model included a scale parameter account for overdispersion of the data through the residual keyword in the RANDOM statement of PROC GLIMMIX in SAS version 9.0. (Stroup, 2013).

### Heritability estimation

The *ad-hoc* broad-sense heritability was estimated as:

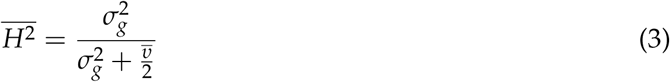

where 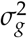 is the genetic variance and 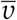 is the mean variance of the difference of the adjusted means (Piepho and Möhring, 2007). To estimate 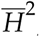, models were fit with genotypes as a random effect using the ASREML-R package for severity and sporulation and PROC GLIMMIX in SAS for incidence to obtain an estimate of 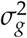. The models with genotypes as fixed effect were used to estimate 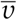.

### Model comparisons

To compare models with different error variance structures, the restricted likelihood ratio test implemented in the asremlPlus R package (Brien, 2019) was used to test if heterogeneous error variances improved the model. The effect of the replicated check varieties on the estimation of the genetic variance was addressed by adding a dummy variable to the severity of infection, sporulation and incidence models (Piepho *et al*., 2006). Such a model was formulated as

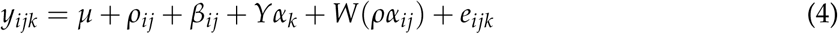

where *Y* and *W* are vectors with 0 for reference varieties and 1 for cultivars and accessions, *α*_*k*_ is the genetic random effect. The remaining effects are the same as in Equations (1) and (2). These models were compared using the mean standard error of the difference (s.e.d.) and heritability. The s.e.d.’s were calculated using the predictplus function of the asremlPlus R package (Brien, 2019).

The effect of transforming our severity of infection and sporulation scorings on heritability estimates was evaluated by repeating the steps outlined above with data transformed with the logit 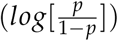 and the angular, or arcsine root, transformation 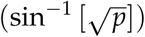, whehre *p* are the severity of infection or sporulation observations. Because the logit function is undefined at 0 or 1, the data at these limits was adjusted by adding and subtracting 0.025 from the original value. The fixed effect of genotypes was tested by using Wald’s F-test as implemented in the ASREML R package for LMMs and type II tests of fixed effects of the Proc GLIMMIX procedure of SAS.

### Comparisons between means

The mean severity and sporulation for the downy mildew infection on each genotype, their confidence intervals and all pairwise comparisons were estimated with the asremlPlus R package (Brien, 2019) for severity and sporulation, and the PROC GLIMMIX procedure of SAS for incidence. Comparisons between the means were based on *t*-tests with a significance threshold *α* = 0.05.

### Correlation between traits

To identify any correlations between phenotypic traits and traits related to the tolerance of quinoa against mildew, i.e. severity, sporulation, and incidence, we used our data from measurements of stomata (width, length, and density). Pearson correlation coefficients were estimated for each pair of variables with a significance threshold *α* = 0.05, using the R package Hmisc (Harrel and Dupont, 2019).

### Relationship between saponin presence and downy mildew severity

Saponin content of seeds was assessed using the foam test (Koziol, 1991). Seeds are placed in a test tube with 5 ml distilled water and shaken vigorously for 30 seconds. Foam height was recorded to the nearest 0.1 cm after shaking. To estimate the robustness of this saponin assay, two seed samples per gene bank accession were evaluated, one from plants grown in Bolivia and one from plants propagated at Højbakkegaard. All accessions with reads equal to zero (i.e., no foam was observed after shaking) were labeled as “no saponin” and all others were marked as “with saponin”. To test for a relationship between saponin presence or absence and downy mildew severity, we conducted a *t*-test using the adjusted means obtained from a LMM with heterogeneous variances between experiments using an untransformed data without the check varieties. This set of means was used because there was no indication from the previous analysis that fitting models with transformed data improved accuracy of the estimates.

### Whole genome DNA sequencing

For DNA extraction, two plants per genotype were grown in a greenhouse of the Taastrup campus at the University of Copenhagen, and two healthy leaves from a single two-months old plant were collected and stored with silica gel for drying. DNA was extracted using the AX Gravity DNA extraction kit (A&A Biotechnology, Gdynia, Poland) following manufacturer’s instructions. Purity and quality of DNA were controlled by agarose gel electrophoresis and concentration determined with a Qubit instrument using SYBR green staining. DNA sequencing libraries were constructed using the protocol of Baym *et al*. (2015). Whole-genome sequencing was done with short-read Illumina sequencing on an Illumina NovaSeq machine (Novogene).

### Genome sequencing, variant calling and genotype imputation

Processing of the raw reads, mapping, and variant calling were done with a custom Snakemake pipeline (Köster and Rahmann, 2012). Raw reads were trimmed with Trim galore v 0.6.4 (Krueger, 2015) (parameters -q 30 --fastqc --paired). Reads were then sorted and indexed with SAMTOOLS 1.10 (Li *et al*., 2009) and deduplicated with the MarkDuplicates (parameter REMOVE DUPLICATES=TRUE tool of PI-CARD v2.21.9 (Broad Institute, Accessed: 2018/02/21; version 2.17.8). The resulting FASTQ files were mapped against the quinoa reference genome version 1.0 (Jarvis *et al*., 2017) and the organellar genomes (Maughan *et al*., 2019) using the Burrows-Wheeler Aligner v0.7.17 (Li and Durbin, 2010) with default parameters.

Variants (SNPs and indels) were called using GATK 3.8 (McKenna *et al*., 2010) by using the HaplotypeCaller tool with a minimum per-base quality score of 20 and a minimum mapping quality score of 30. The GVCF files per sample were merged with the GenotypeGVCFs tool of GATK with default parameters. Missing data was imputed and filtered using LinkImputeR 1.2.3, which allows the user to define a series of filters and evaluate their effect on the accuracy and final number of imputed markers (Money *et al*., 2017). Thresholds for imputation were depth = 8 (Number of reads including a position) and missingness = 0.7 (Proportion of positions/samples with less than the threshold depth). Variant data were filtered to a minor allele frequency *>*= 0.05 and a deviation from Hardy-Weinberg equilibrium *p >*0.01 using a likelihood ratio test (Maruki and Lynch, 2015). Linkage disequilibrium was estimated with a pair-wise correlation coefficient between variants, *r*^2^, using the final VCF file as input for PopLDdecay (Zhang *et al*., 2018) with default parameters.

### Genome-wide association study (GWAS)

We used two methods for association mapping, Farm-CPU, for use with sequence variants (SNPs) and a *k*-mer-based method. FarmCPU (Fixed and random model Circulating Probability Unification) uses SNP in a two-step iterative process with fixed and random effects models to improve computation times, reduce the confounding effects of structure and improve power to identify significant marker-trait associations in comparison to other methods (Liu *et al*., 2016; Malik *et al*., 2019). The model was run with and without the inclusion of the first three principal components with a *p*-value threshold of 0.01 (Bonferroni corrected) for both the inclusion of a marker during the first iteration of the model as well as the genome-wide significance threshold.

The *k*-mer based method by Voichek and Weigel (2020) identifies genotype-phenotype associations using sequencing reads instead of molecular variants to address the lack of a reference genome or account for structural variation. We implemented the method in a Snakemake pipeline using the following parameters: *k*-mer length of = 31 nucleotides, minor allele count = 3 minor allele frequency = 0.05. This method requires a kinship matrix, which was estimated with a method used by EMMA (Efficient Mixed-Model Association) and consists of an identical-by-state (IBS) allele-sharing matrix under the assumption that every variant has a small random effect on the phenotype (Kang *et al*., 2008).

## Acknowledgments

We are grateful to INIAF (Bolivian National Institute for Innovation in Agriculture and Forestry) and to the greenhouse team. The authors wish to thank laboratory technician Lene Klem at University of Copenhagen for the PCR analysis on the pathogen. We thank Elisabeth Kokai-Kota for the construction of DNA sequencing libraries and to Mireia Vidal for advice on bioinformatics. The authors acknowledge support by the state of Baden-Württemberg through bwHPC. We express our appreciation to Sven-Eric Jacobsen for the provision of the Danish cultivars, support and knowledge.

## Declarations

### Ethics approval and consent to participate

Not applicable.

### Consent for publication

Not applicable.

### Availability of data and materials

All phenotypic data generated or analysed during this study are included in this published article in Supplementary File 1 (Complete mildew infection dataset). The sequencing data for the identification of *Peronospora variabilis* are available from NCBI Genebank with ID MT895880. Raw whole genome sequencing read data are available from the European Nucleotide Archive (ENA) with project number ID PRJEB39907 (Accessions ERS4955326-ERS4955413). Analysis scripts are available from Zenodo archive with DOI:

## Authors contributions

CCL, MCA, HPP and KS designed the study. CCL conducted the field house experiment and phenotyping work. OSL designed the pilot study and the molecular identification of the pathogen isolate. CCL and MCA analysed the phenotyping data. DBA, CA and SS advised on experimental work and data analysis. MCA analysed sequencing data and carried out GWAS. CCL, MCA and KS wrote the manuscript. All authors read, revised and agreed on the manuscript.

## Additional Files

- Additional File 1: Supplementary Figures S1 to S5 and Supplementary Tables S1 to S3.
- Additional File 2: Supplementary File 1: List of genebank accessions and passport data; Mildew infection raw data; saponin and stomatal measurements.
- Additional File 3: Supplementary File 2: Detailed protocol for the isolation and maintenance of the downy mildew pathogen *Peronospora variabilis*.

