## Supplementary File 2 for "Genetic variation for tolerance to the downy mildew pathogen *Peronospora variabilis* in genetic resources of quinoa (*Chenopodium quinoa*)"

### Protocols of the experimental design

- List of biological control agents used in this study

Table 1: Species used to control pests

| *Species* |
| --- |
| *Aphidius Col/Erv/Aphel* |
| *Macrolophus caliginosus* |
| *Amblyseius cucumeris* |
| *Aphidoletes aphidimyza* |
| *Orius majusculus/laevigatus* |
| *Encarsia Formosa* |
| *Hypoaspis Miles* |
| *Amblyseius swirskii* |
| *Amblyseius Californicus* |
| *Chrysoperla carnea* |

- Protocol for propagation of the *Peronospora variabilis* isolate
  - Rinse infected leaves with distilled sterile water.
  - Filter the solution and re-suspend the spores in sterile water.
  - Calibrate the spore's concentration to $10^5$ spores per ml; give the solution a cold snap for 5 minutes; load the bottle feeder of an air brush.
  - Set the pressure to 50 bar and spray 7 to 8 weeks old potted quinoa plants of the Danish cultivar Vikinga and Bolivian cultivar Blanca, making sure to cover all the leaves
  - Cover inoculated plants with polyethylene bags and keep them in a dark chamber at 22degree C during the day and 15degree C during the night for 48 hours.
  - Remove the bags and keep the plants under a 16h photoperiod and a 22degree C/18degree C day/night temperature cycle for five days, until the first signs of infection are visible.
  - Cover the plants with polyethylene bags again, reduce nighttime temperature to 15degree C, and keep them in a dark chamber for 24 hours. At this point, there should be signs of sporulation on the abaxial side of the leaves.
  - Collect leaves with signs of sporulation and prepare a spore solution.
  - Repeat this process as necessary.
