## Supplementary Tables and Figures for "Genetic variation for tolerance to the downy mildew pathogen *Peronospora variabilis* in genetic resources of quinoa (*Chenopodium quinoa*)"

### Supplementary material 1

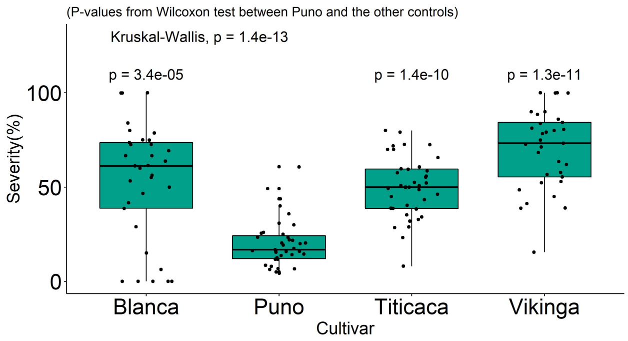


Supplementary Figure S1: Boxplot of the control genotypes from the pilot experiment

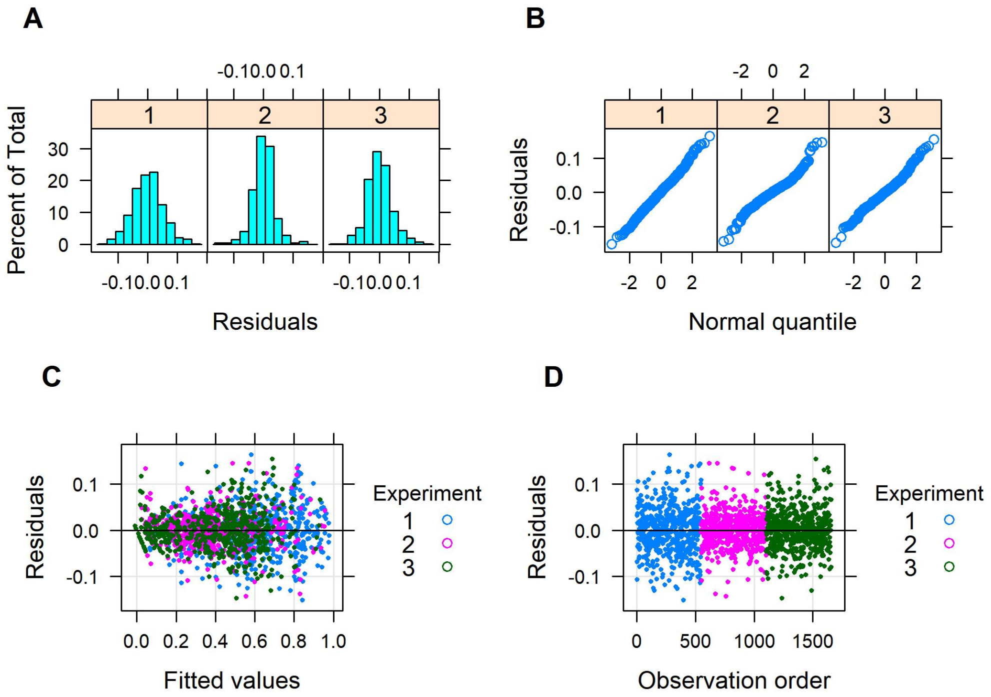


Supplementary Figure S2: Diagnostic plots for a severity LMM with untransformed data and checks included. A) histogram of residuals. B) qq-plot. C) Fitted vs. Residuals plot. D) plot of ordered residuals by experiment.

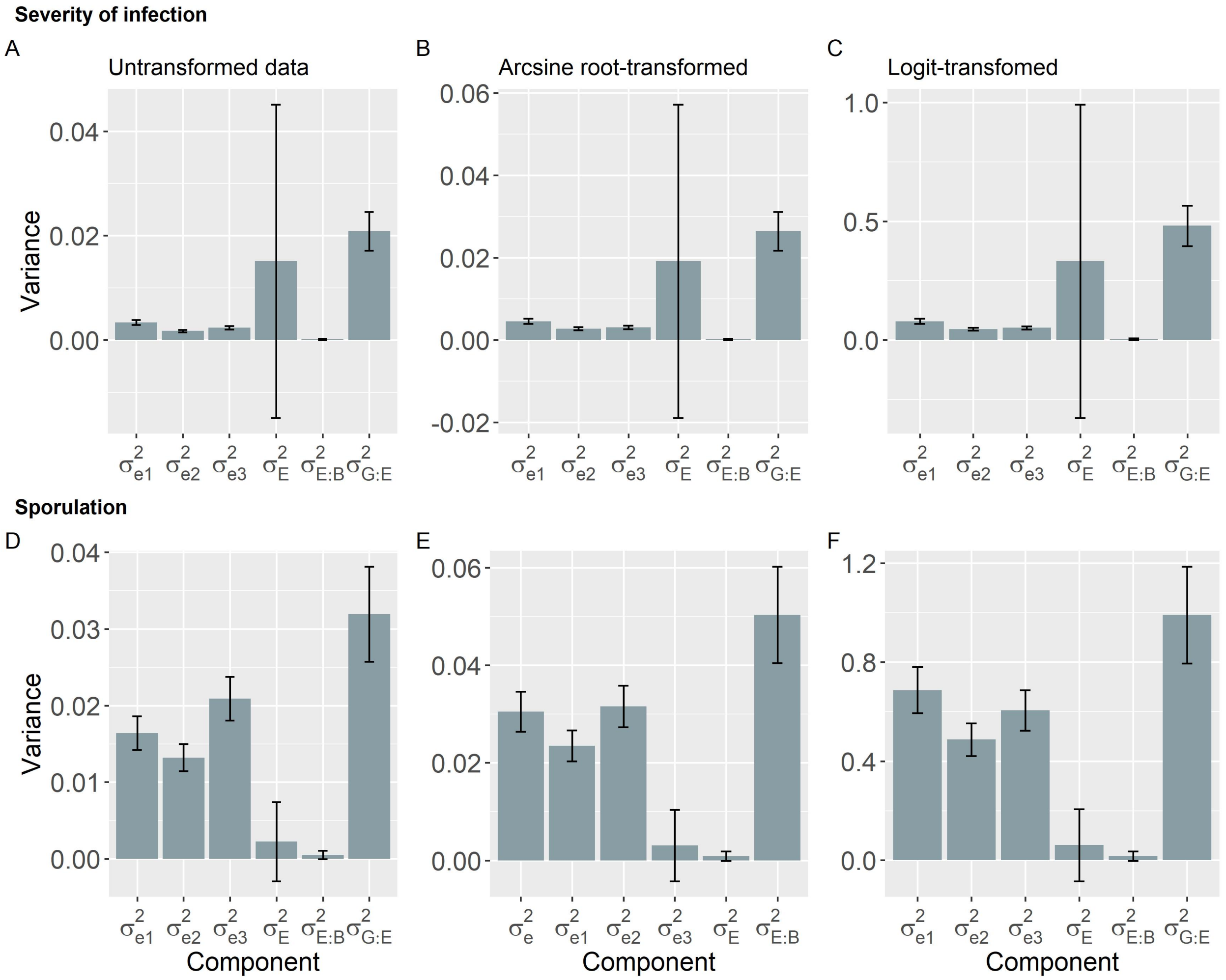


Supplementary Figure S3: Variance components for the trait severity of infection (A-C) and Sporulation (D-F) as estimated with a linear mixed model including the checks. Models were fitted (A,D) without data transformation, (B,E) Arcsine root transformation, and (C,F) logit transformation. $\sigma^2_{e1}$, $\sigma^2_{e2}$, $\sigma^2_{e3}$ are residual variances for experiments 1-3; $\sigma^2_{E}$; $\sigma^2_{E:B}$ and $\sigma^2_{G:E}$ are variance components for experiments, blocks nested within experiments and the genotype by experiment interaction, respectively.

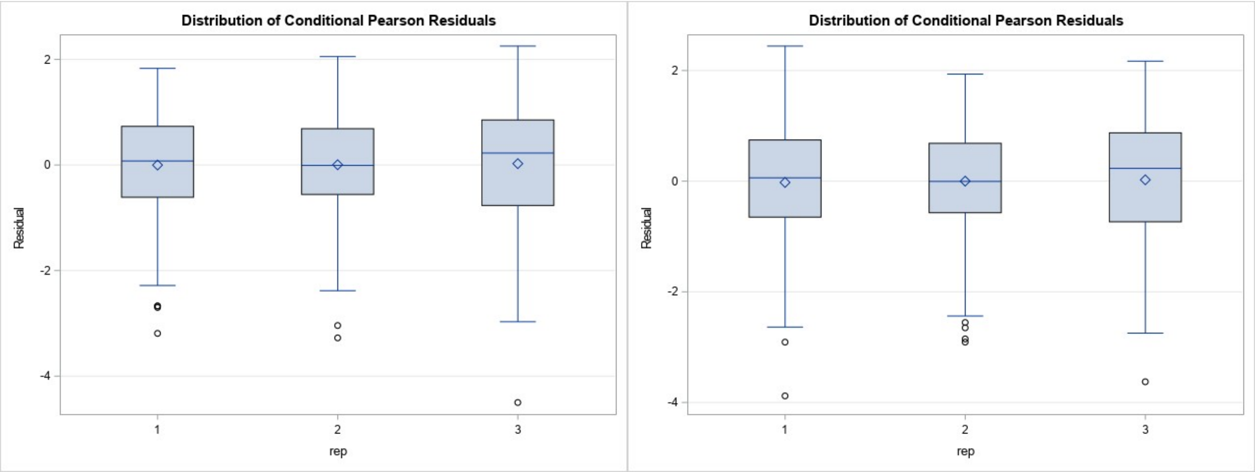


Supplementary Figure S4: Boxplot of the Pearson conditional residuals of the incidence GLMM with (left) and without (right) checks.

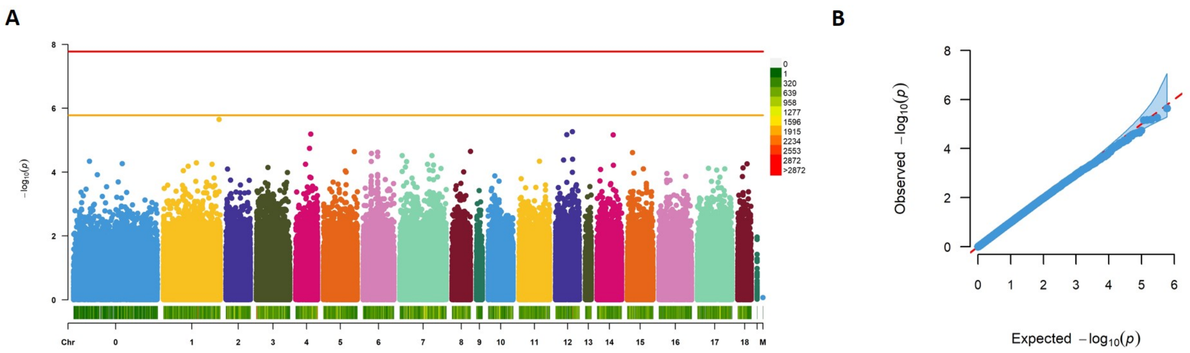


Supplementary Figure S5: Association mapping for downy mildew severity using FarmCPU with models with 3 principal components as covariates. Manhattan plot (A). Red line shows the Bonferroni corrected threshold for $p = 0.01$ and orange line indicates a suggestive threshold (1/number of markers). Bar at the bottom indicates marker density. QQ plot (B) for the FarmCPU model with the 95% confidence interval (light blue); Red line draws the expected distribution of $p$-values.

Supplementary Table S1: Results of the Restricted Likelihood Ratio Test for the disease severity and sporulation of downy mildew

| *Variable* | *Model* | *Test statistic* | *DF* | *p-value* |
|---|---|---|---|---|
| *Severity* | Untransformed - With Checks | 44.965 | 2 | $1.72x10^{-10}$ |
| *Severity* | Untransformed - No Checks | 17.119 | 2 | $1.92x10^{-04}$ |
| *Severity* | Arcsine root - With Checks | 29.151 | 2 | $4.68x10^{-07}$ |
| *Severity* | Arcsine root - No Checks | 13.106 | 2 | $1.43x10^{-03}$ |
| *Severity* | Logit - With Checks | 35.056 | 2 | $2.44x10^{-08}$ |
| *Severity* | Logit - No Checks | 23.072 | 2 | $9.77x10^{-06}$ |
| *Sporulation* | Untransformed - With Checks | 22.276 | 2 | $1.45x10^{-05}$ |
| *Sporulation* | Untransformed - No Checks | 6.486 | 2 | $3.90x10^{-02}$ |
| *Sporulation* | Arcsine root - With Checks | 10.646 | 2 | $4.88x10^{-03}$ |
| *Sporulation* | Arcsine root - No Checks | 3.219 | 2 | 0.199 |
| *Sporulation* | Logit - With Checks | 12.579 | 2 | $1.86x10^{-03}$ |
| *Sporulation* | Logit - No Checks | 7.307 | 2 | $2.59x10^{-02}$ |

(Note: each line is a test comparing a null model with homogeneous variances for the error with a model with a heterogeneous variance structure.)

Supplementary Table S2: Estimated variance components for the models fitted for severity and limits for the 95% confidence intervals

| Model | Variance Component | Estimate | 95% CI lower limit | 95% CI upper limit |
|---|---|---|---|---|
| Untransformed - No Checks | Experiment:Block | 0.0001 | $-1.14 \times 10^{-05}$ | $2.55 \times 10^{-04}$ |
| Untransformed - No Checks | Gen:Exp | 0.0225 | $1.84 \times 10^{-02}$ | $2.66 \times 10^{-02}$ |
| Untransformed - No Checks | Exp-1 Error Variance | 0.0036 | $3.14 \times 10^{-03}$ | $4.13 \times 10^{-03}$ |
| Untransformed - No Checks | Exp-2 Error Variance | 0.0023 | $1.98 \times 10^{-03}$ | $2.65 \times 10^{-03}$ |
| Untransformed - No Checks | Exp-3 Error Variance | 0.0032 | $2.72 \times 10^{-03}$ | $3.62 \times 10^{-03}$ |
| Arcsine root - With Checks | Experiment | 0.0192 | $-1.89 \times 10^{-02}$ | $5.72x10^{-02}$ |
| Arcsine root - With Checks | Experiment:Block | 0.0002 | $-9.63 \times 10^{-06}$ | $3.84 \times 10^{-04}$ |
| Arcsine root - With Checks | Gen:Exp | 0.0264 | $2.17 \times 10^{-02}$ | $3.11 \times 10^{-02}$ |
| Arcsine root - With Checks | Exp-1 Error Variance | 0.0046 | $3.93 \times 10^{-03}$ | $5.19 \times 10^{-03}$ |
| Arcsine root - With Checks | Exp-2 Error Variance | 0.0028 | $2.39 \times 10^{-03}$ | $3.14 \times 10^{-03}$ |
| Arcsine root - With Checks | Exp-3 Error Variance | 0.0031 | $2.67 \times 10^{-03}$ | $3.50 \times 10^{-03}$ |
| Arcsine root - No Checks | Experiment | 0.0239 | $-2.31 \times 10^{-02}$ | $7.09 \times 10^{-02}$ |
| Arcsine root - No Checks | Experiment:Block | 0.0002 | $-1.80 \times 10^{-05}$ | $3.33 \times 10^{-04}$ |
| Arcsine root - No Checks | Gen:Exp | 0.0276 | $2.26 \times 10^{-02}$ | $3.27 \times 10^{-02}$ |
| Arcsine root - No Checks | Exp-1 Error Variance | 0.0047 | $4.05 \times 10^{-03}$ | $5.35 \times 10^{-03}$ |
| Arcsine root - No Checks | Exp-2 Error Variance | 0.0034 | $2.89 \times 10^{-03}$ | $3.85 \times 10^{-03}$ |
| Arcsine root - No Checks | Exp-3 Error Variance | 0.005 | $4.29 \times 10^{-03}$ | $5.74 \times 10^{-03}$ |
| Logit - With Checks | Experiment | 0.3318 | $-3.28 \times 10^{-01}$ | $9.91 \times 10^{-01}$ |
| Logit - With Checks | Experiment:Block | 0.0033 | $-1.64 \times 10^{-04}$ | $6.76 \times 10^{-03}$ |
| Logit - With Checks | Gen:Exp | 0.4813 | $3.96 \times 10^{-01}$ | $5.67 \times 10^{-01}$ |
| Logit - With Checks | Exp-1 Error Variance | 0.0788 | $6.78 \times 10^{-02}$ | $8.97 \times 10^{-02}$ |
| Logit - With Checks | Exp-2 Error Variance | 0.0455 | $3.93 \times 10^{-02}$ | $5.17 \times 10^{-02}$ |
| Logit - With Checks | Exp-3 Error Variance | 0.0512 | $4.42 \times 10^{-02}$ | $5.82 \times 10^{-02}$ |
| Logit - No Checks | Experiment | 0.4876 | $-4.72 \times 10^{-01}$ | 1.45 |
| Logit - No Checks | Experiment:Block | 0.0031 | $-3.06 \times 10^{-04}$ | $6.46 \times 10^{-03}$ |
| Logit - No Checks | Gen:Exp | 0.5059 | $4.13 \times 10^{-01}$ | $5.99 \times 10^{-01}$ |
| Logit - No Checks | Exp-1 Error Variance | 0.0838 | $7.20 \times 10^{-02}$ | $9.55 \times 10^{-02}$ |
| Logit - No Checks | Exp-2 Error Variance | 0.0549 | $4.71 \times 10^{-02}$ | $6.27 \times 10^{-02}$ |
| Logit - No Checks | Exp-3 Error Variance | 0.0944 | $8.07 \times 10^{-02}$ | $1.08 \times 10^{-01}$ |

Supplementary Table S3: Estimated variance components for the models fitted for sporulation and limits for the 95% confidence intervals

| **Model** | **Variance Component** | **Estimate** | **95% CI lower limit** | **95% CI upper limit** |
|---|---|---|---|---|
| Untransformed - With Checks | Experiment | 0.0022 | $-2.92 \times 10^{-03}$ | $7.38 \times 10^{-03}$ |
| Untransformed - With Checks | Experiment:Block | 0.0005 | $-6.68 \times 10^{-05}$ | $1.03 \times 10^{-03}$ |
| Untransformed - With Checks | Gen:Exp | 0.0319 | $2.57 \times 10^{-02}$ | $3.82 \times 10^{-02}$ |
| Untransformed - With Checks | Exp-1 Error Variance | 0.0164 | $1.42 \times 10^{-02}$ | $1.86 \times 10^{-02}$ |
| Untransformed - With Checks | Exp-2 Error Variance | 0.0132 | $1.14 \times 10^{-02}$ | $1.50 \times 10^{-02}$ |
| Untransformed - With Checks | Exp-3 Error Variance | 0.0209 | $1.81 \times 10^{-02}$ | $2.38 \times 10^{-02}$ |
| Untransformed - No Checks | Experiment | 0.0093 | $-9.51 \times 10^{-03}$ | $2.81 \times 10^{-02}$ |
| Untransformed - No Checks | Experiment:Block | 0.0005 | $-8.63 \times 10^{-05}$ | $1.01 \times 10^{-03}$ |
| Untransformed - No Checks | Gen:Exp | 0.0365 | $2.90 \times 10^{-02}$ | $4.40 \times 10^{-02}$ |
| Untransformed - No Checks | Exp-1 Error Variance | 0.0188 | $1.62 \times 10^{-02}$ | $2.14 \times 10^{-02}$ |
| Untransformed - No Checks | Exp-2 Error Variance | 0.0166 | $1.43 \times 10^{-02}$ | $1.90 \times 10^{-02}$ |
| Untransformed - No Checks | Exp-3 Error Variance | 0.0216 | $1.87 \times 10^{-02}$ | $2.46 \times 10^{-02}$ |
| Arcsine root - With Checks | Experiment | 0.0031 | $-4.24 \times 10^{-03}$ | $1.04 \times 10^{-02}$ |
| Arcsine root - With Checks | Experiment:Block | 0.0009 | $-1.13 \times 10^{-04}$ | $1.84 \times 10^{-03}$ |
| Arcsine root - With Checks | Gen:Exp | 0.0503 | $4.05 \times 10^{-02}$ | $6.02 \times 10^{-02}$ |
| Arcsine root - With Checks | Exp-1 Error Variance | 0.0305 | $2.64 \times 10^{-02}$ | $3.46 \times 10^{-02}$ |
| Arcsine root - With Checks | Exp-2 Error Variance | 0.0235 | $2.03 \times 10^{-02}$ | $2.67 \times 10^{-02}$ |
| Arcsine root - With Checks | Exp-3 Error Variance | 0.0316 | $2.73 \times 10^{-02}$ | $3.58 \times 10^{-02}$ |
| Arcsine root - No Checks | Experiment | 0.0094 | $-1.00 \times 10^{-02}$ | $2.89 \times 10^{-02}$ |
| Arcsine root - No Checks | Experiment:Block | 0.0009 | $-1.41 \times 10^{-04}$ | $1.87 \times 10^{-03}$ |
| Arcsine root - No Checks | Gen:Exp | 0.0542 | $4.31 \times 10^{-02}$ | $6.53 \times 10^{-02}$ |
| Arcsine root - No Checks | Error Variance | 0.031 | $2.86 \times 10^{-02}$ | $3.34x10^{-02}$ |
| Logit - With Checks | Experiment | 0.0612 | $-8.42 \times 10^{-02}$ | 0.207 |
| Logit - With Checks | Experiment:Block | 0.0168 | $-2.48 \times 10^{-03}$ | $3.60 \times 10^{-02}$ |
| Logit - With Checks | Gen:Exp | 0.9906 | 0.795 | 1.19 |
| Logit - With Checks | Exp-1 Error Variance | 0.6876 | 0.595 | 0.780 |
| Logit - With Checks | Exp-2 Error Variance | 0.4873 | 0.422 | 0.553 |
| Logit - With Checks | Exp-3 Error Variance | 0.6055 | 0.523 | 0.688 |
| Logit - No Checks | Experiment | 0.1829 | $-0.195$ | 0.561 |
| Logit - No Checks | Experiment:Block | 0.0172 | $-2.83 \times 10^{-03}$ | $3.73 \times 10^{-02}$ |
| Logit - No Checks | Gen:Exp | 1.0652 | 0.846 | 1.28 |
| Logit - No Checks | Exp-1 Error Variance | 0.7415 | 0.64 | 0.843 |
| Logit - No Checks | Exp-2 Error Variance | 0.5657 | 0.487 | 0.644 |
| Logit - No Checks | Exp-3 Error Variance | 0.6145 | 0.531 | 0.698 |
